# Molecular Dynamics Simulations on Quercetin-3-(6-Malonylglucoside) From *Morus Alba* Shows Optimal Inhibition of Bcl-2 with Favorable Anti-Tumor Activities

**DOI:** 10.1101/2023.07.04.547659

**Authors:** Emmanuel Sunday Omirin, Olaposi Idowu Omotuyi, Oluwaseun Grace Afokhume, Ehisdiame Favour Okoh, Samuel Oluwaseun Boboye, Babatunde Oluwaseun Ibitoye, Olabode Oluwagbemiga Adelegan, Ezekiel Abiola Olugbogi, Michael Aladejare Aderiye, Oluwafemi Ojo Agosile

## Abstract

The target of most cancer chemotherapeutic agents is to drive cancer cells toward death. A fine balance between anti-apoptotic and pro-apoptotic proteins is needed to maintain cellular homeostasis. Any shift favoring the pro-apoptotic ones is needed to drive cellular death in cancer chemotherapy. However, anti-apoptotic proteins such as Bcl-2 and Bcl-xL bind with pro-apoptotic proteins to hinder apoptosis mechanisms. Overexpression of these anti-apoptotic proteins lead to several cancers by preventing apoptosis. In this study, molecular docking, ADMET predictions, and molecular dynamics simulations were performed for the identification of potent inhibitors of anti-apoptotic Bcl-2 with compounds of *Morus alba.* Our study discovered that Quercetin-3-(6-Malonylglucoside) and Epigallocatechin gallate recorded excellent binding affinity with Bcl-2. Therefore, we conclude that compounds of *Morus alba* should be subjected to further experimental studies (*in vitro* and *in vivo)* in order to confirm the findings that they could be used as better options in cancer chemotherapy.

## 1.0 Introduction

The anti-apoptotic Bcl-2 proteins, such as Bcl-2 and Bcl-xL, are fast becoming important targets in the fight against many forms of known cancer. Bcl-2, found in the outer membrane of the mitochondria, is an important protein that plays important roles in the maintenance of cellular homeostasis, a balance between cell death and cell survival (Kelly and Strasser, 2020). It is a member of the Bcl-2 family of proteins. This family of proteins consist of members that either promote or hinder apoptosis, and thus mediate apoptosis by taking control of the mitochondrial outer membrane permeabilization (MOMP), a key step in the intrinsic arm of apoptosis (Youle and Strasser, 2008). According to Chao and Korsmeyer (1998), the members of the family involved in pro-apoptosis include but not limited to Bax, Bak, Diva, Bcl-xS, Bik and Bim; while those with anti-apoptosis include Bcl-2, Bcl-xL, Mcl-1 and Bfl-1. In cancer, a disturbance in the homeostatic balance between cell growth and cell death, Bcl-2 has registered its presence as one of the major players, for it has been identified as a cause of a number of cancers, including melanoma, breast, prostate, chronic lymphocytic leukemia, and lung cancer, and a possible cause of schizophrenia and autoimmunity (Garcia-Aranda et al., 2018).

Since the roles of Bcl-2 have been established in cancer, there is an important need to explore various ways of controlling the protein, being an anti-apoptotic protein. Various mechanisms could be used to regulate the activities of this protein (Zhou et al., 2011). These mechanisms include gene expression regulation which requires that its expression be tightly regulated at transcriptional level, for instance, the inhibition of the PI3K/AKT pathway, which promote cell survival, can downregulate Bcl-2 expression (Hildeman et al., 2003); post-transcriptional regulation which uses several regulatory mechanisms to control the abundance and stability of the mRNA of Bcl-2 which modulates its translation and degradation (Bevilacqua et al., 2003; Chen et al., 2016); post-translational modifications, phosphorylation by kinase for example, can regulate its anti-apoptotic function. Conversely, pro-apoptotic signals can induce proteasomal degradation of Bcl-2, leading to reduced levels of the protein and enhanced apoptosis (Czabotar et al., 2014; Llambi and Green, 2011); protein-protein interactions, epigenetic regulation, and protein-ligand interactions which involve the use of small molecules as the antagonists of Bcl-2 protein. Inhibition of Bcl-2 in cancer has emerged as a promising therapeutic approach, particularly in hematological malignancies (He and Hu, 2018). These inhibitors are made to target and counteract the anti-apoptotic function of Bcl-2, thus promoting the death of cancerous cells.

In the present study, *in silico* approaches using pharmacokinetic and pharmacodynamics properties, molecular docking, and molecular dynamics (MD) simulations were used to show that compounds from *Morus alba* are capable of inhibiting Bcl-2, and that the inhibition of Bcl-2 is needed to drive cellular death in actively dividing and cancerous cells.

### 1.1 Morus alba

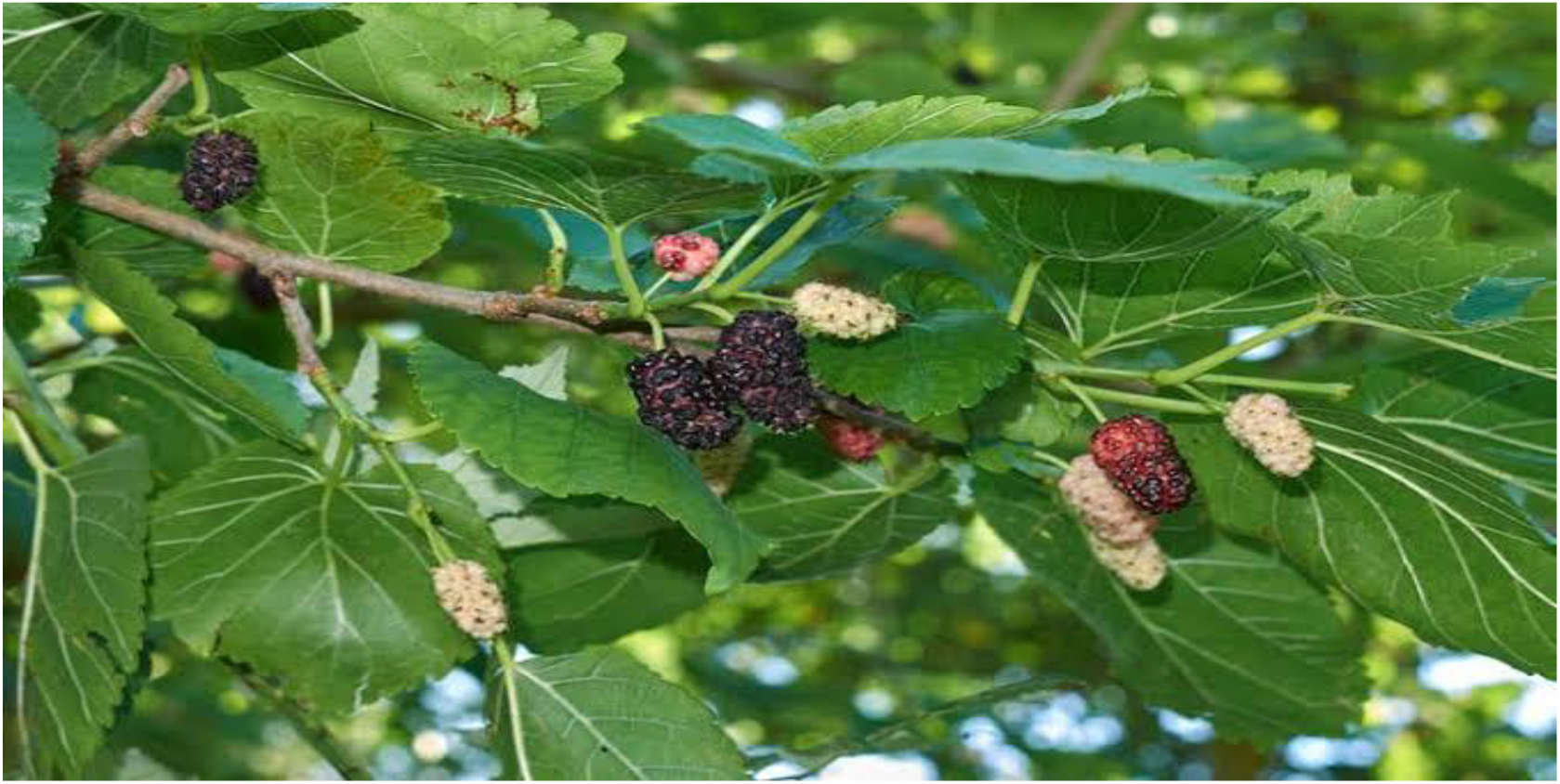

Historically, plants have been used since ancient times in traditional communities for the treatment of many diseases (WHO, 2008). *Morus alba* (mulberry) has a long history of use as fodder and traditional medicine. There has been much work done and published on this species (Chan et al., 2016). Pharmacologically, the plant has been reported to have antioxidant property, due to being a rich source of anthocyanins, compounds which are excellent antioxidant agent with strong free radical scavenging potency than known standards like vitamin C (Bae and Suh, 2007). Similarly, Kuwanon G isolated from the methanol root back extract of the plant demonstrated antimicrobial activity against pathogens such as *Streptococcus mutans, Streptococcus sanguis and Staphylococcus sobrinus* as reported by Park and colleagues (2003). In similar manner, the cytotoxic effect of the plant has been tested by Kofujita and co. (2004) where a flavone compound (7, 2′, 4′, 6′-tetrahydoroxy-6-geranylflavanone) isolated from the plant inhibited the growth of rat hepatoma in dRLh84 cells with an IC^50^ value of 53microgram per mole. Overall, two flavonioids (quercetin-3-O-β-D-glucopyranoside and quercetin-3-7-di-O-β-D-glucopyranoside) found with the plant were found to inhibit the growth of human leukemia HL-60 cells (Kim et al., 2000). Therefore, the work presented herein was inspired by the reported therapeutic effects and safety of the plant, and a need to discover new natural blockers of Bcl-2 that could be better employed in the long fight against human cancer.

### 1.2 Bcl-2 Homology Domain 3 (BH3) Mimetic as Bcl-2 inhibitors

Since the anti-apoptotic Bcl-2 proteins, such as Bcl-2 and Bcl-xL, conserve all four BH domains (Reed et al., 1996), seeking to use a typeable Bcl-2 homology (BH) mimetic in controlling the activities of the protein represents an important area of the fight against cancer. BH3 mimetics are a class of small molecule compounds that have garnered significant attention as potent inhibitors of Bcl-2 family proteins, particularly Bcl-2 itself (Yap et al., 2017). These inhibitors mimic the function of the BH3 domain, a critical region found in pro-apoptotic proteins of the Bcl-2 family, enabling them to selectively bind and neutralize the anti-apoptotic activities of Bcl-2 (Gross et al., 1999). The development of BH3 mimetics represents a promising strategy to induce apoptosis in cancer cells, making them attractive candidates for targeted cancer therapy (Sarosiek and Letai, 2016; Labi et al., 2008). A lot of these compounds have been approved for their chemotherapeutic use. Very popular among these compounds are ABT-199 (Venetoclax) and ABT-263 (Navitoclax).

**Figure 2.**
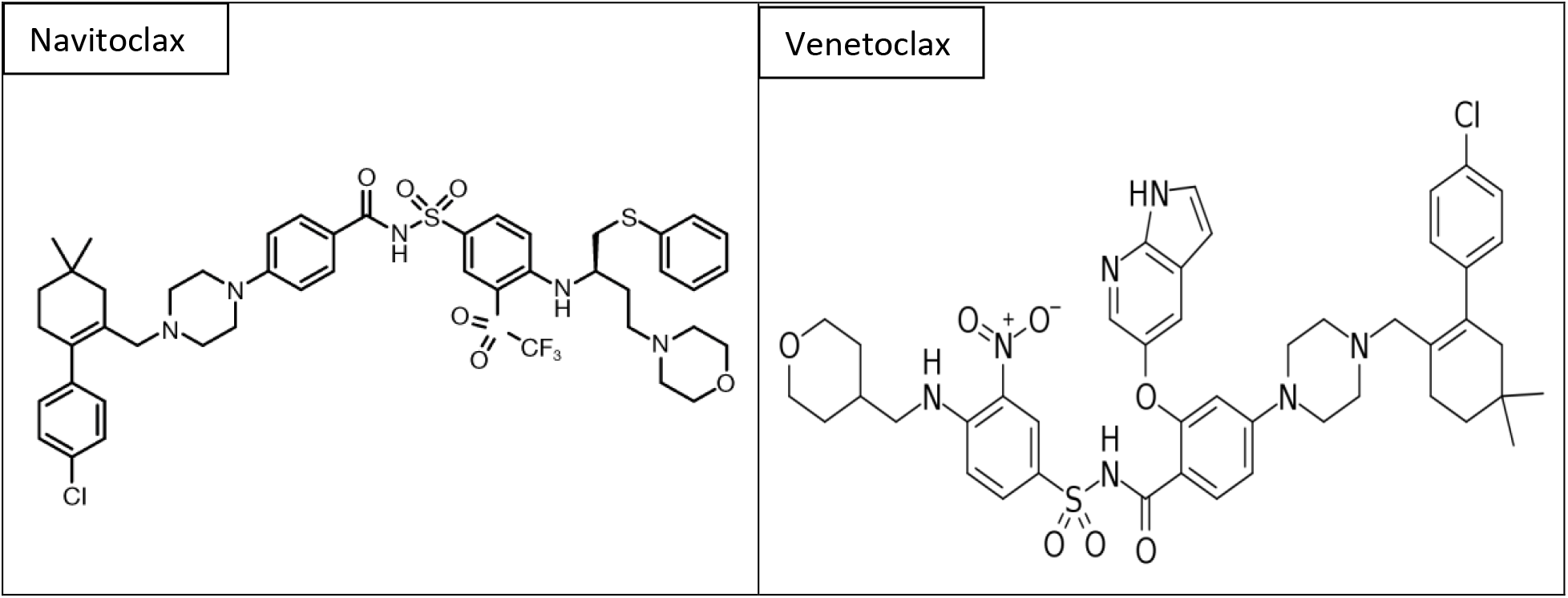
Chemical structures of navitoclax (left) and venetoclax (right) as retrieved from Wikipedia.

The development of targeted therapies have shed more light on the field of oncology, providing new treatment options for various cancers. Venetoclax has received approval for several indications, including chronic lymphocytic leukemia (CLL), and acute myeloid leukemia (AML) by selectively inhibiting Bcl-2 and thus arrest the growth of cancerous cells. On the other hand, navitoclax has demonstrated efficacy in indications including CCL, myelofibrosis, and other blood related malignancies. Though it has been established that these chemotherapeutic agents often come with some deleterious effects, Souers and colleagues (2013) demonstrated that Venetoclax is capable of inhibiting the growth of Bcl-2-dependent tumors in vivo and also spare human platelets. These researchers concluded that a single dose of ABT-199 in three patients with refractory chronic lymphocytic leukemia resulted in tumor lysis within 24 hours.

## 2.0 Computational Approaches

All computational studies which include ligand preparation, protein preparation, receptor grid generation, molecular docking, MM-GBSA and aspects of ADMET predictions were done with the various modules available in Schrodinger Maestro software version 12.8.

### 2.1 Target identification, preparation, and receptor grid generation

The structure of human B-cell lymphoma 2 (Bcl-2) in complex with Venetoclax (ABT-199) was downloaded from the RCSB protein data bank (https://www.rcsb.org/structure/4MAN) (Souers et al., 2013). The choice of the target was informed by the need to produce identical but better result than what has been reported. The retrieved protein was uploaded to Schrodinger Maestro and prepared. The preparation involves pre-processing, addition of bond orders, hydrogen atoms, filling loops, and removing water molecules beyond 5.00Å. The processed protein was subjected to interactive optimization to refine the crystallized protein structure and restrained minimization converging heavy atoms to RMSD at 0.30Å.

**Figure 3:**
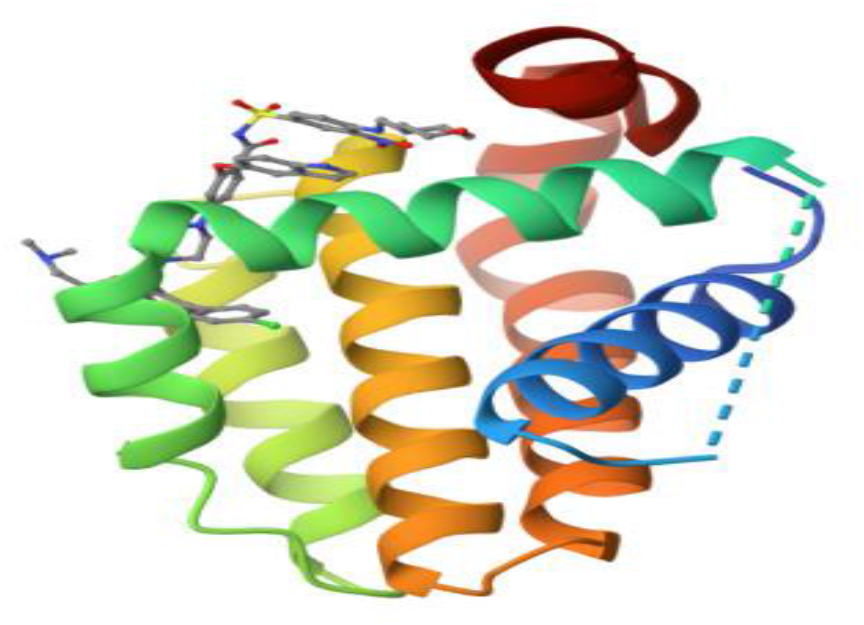
Structure of the human Bcl-2 (4MAN) in complex with venetoclax (ABT-199) as deposited by Souers et al., 2013.

Receptor grid generation defines the binding orientation and the size of the active site for protein-ligand docking. The receptor grid was generated based on the co-crystallized ABT-199 ligand present in the target protein. The grid area is determined using the receptor grid generation feature to identify the region in the system that serves as a receptor. The grid is set with the inhibitory center of ABT-199, the native ligand of PDB ID 4MAN, at the protein’s active site forming XYZ center coordinates of -11.72, 9.94, and 9.02 respectively.

### 2.2 Ligand preparation, Lipinski’s rule (LRoV) and QikProp screening of compounds

A library of ninety-one (91) selected phytochemicals reported in *Morus alba* by Chan et al., 2016 were retrieved from PubChem database (https://pubchem.ncbi.nlm.nih.gov). The ligands, during the preparation, were neutralized and just one stereoisomer was generated at most for all ligands using the LigPrep module in Schrodinger Maestro (LigPrep, 2018). After this, they were subjected to QikProp to screen compounds with druglike characters. A lot of these compounds were fit for drug development; they were further subjected to the pharmacophore hypothesis, to check which compounds with matching features of crystallized protein-ligand complex.

### 2.3 Molecular docking

Glide-XP (extra precision) was used to dock the prepared ligands into the designated active site of the prepared protein guided by the grid generated. The Van Der Waals scaling factor was set at 0.80 for the ligands atoms.

### 2.4 Binding free energy calculation/thermodynamics calculation

To determine the binding free energy of the docked complexes, the molecular mechanics with generalized born surface area (MM-GBSA) tool integrated with prime of the Schrödinger Maestro 12.8 was used.

### 2.5 Pharmacophore hypothesis, ligand screening, and ADMET prediction

Energy-optimized pharmacophore hypothesis was generated using the crystal structure of 4MAN linked to its co-crystal ligand, ABT-199. The E-pharmacophore model was created using the phase module develop pharmacophore from protein-ligand complex option. The auto-pharmacophore approach was used. For the hypothesis settings, features that made interactions with the protein were chosen, and then a receptor-based excluded volume shell was created to mimic the receptor binding site, ignoring receptor atoms whose surfaces are within 2.00 Å of the ligand surface, and limiting excluded volume shell thickness to 5.00 Å. E-pharmacophore-based virtual screening was performed using Maestro Schrodinger 2018, version 12.8 according to the method reported by Omoboyowa and colleagues (2022).

Toxicological predictions of some of the ligands were carried out using the QikProp modules of Maestro Schrodinger to see if the ligands were safe if used as human drugs. Later, they were further screened using SwissADME (http://www.swissadme.ch/index.php) and Protox II (https://tox-new.charite.de/protox_II/index.php?site=compound_input). Prediction was made by writing the canonical smiles string of the ligand compound and then selecting what properties are to be predicted for example absorption (water solubility, intestinal absorption, and skin permeability) distribution, metabolism, excretion, and toxicity.

### 2.6 Molecular dynamics simulation

Docked complexes of the hit compound, Quercetin-3-(6-Malonylglucoside), the second top-ranked compound, Epigallocatechin gallate, and the standard drugs (Navitoclax and Venetoclax) were subjected to molecular dynamics (MD) simulation studies using the Desmond module of Maestro Schrodinger. The primary objective of this simulation was to assess the stability of the complexes and validate the docking results obtained earlier. During the simulation, the complexes were allowed to undergo a 100ns simulation using the NPT ensemble class, at a constant temperature of 300.0K and pressure of 1.01325 bar. The system was prepared using the System Builder module, which employed the TIP3P solvent model. An orthorhombic boundary box with dimensions of 10 x 10 x 10 Å was used, and the OPLS3e force field was employed (Balogun et al., 2022). The box was minimized, and the system charges were neutralized by the addition of Na^+^ and Cl^-^ ions. To monitor the stability of the ligands and protein in their native motion, root mean square deviation (RMSD) and root mean square fluctuation (RMSF) were estimated. RMSD was used to measure the deviation of the simulated system from a reference structure, while RMSF quantified the fluctuation of each atom in the system during the simulation. The analysis of RMSD and RMSF provided a detailed understanding of the conformational dynamics and stability of the simulated complexes.

## 3.0 Results

### 3.1 Pharmacophore hypothesis

**Figure 4:**
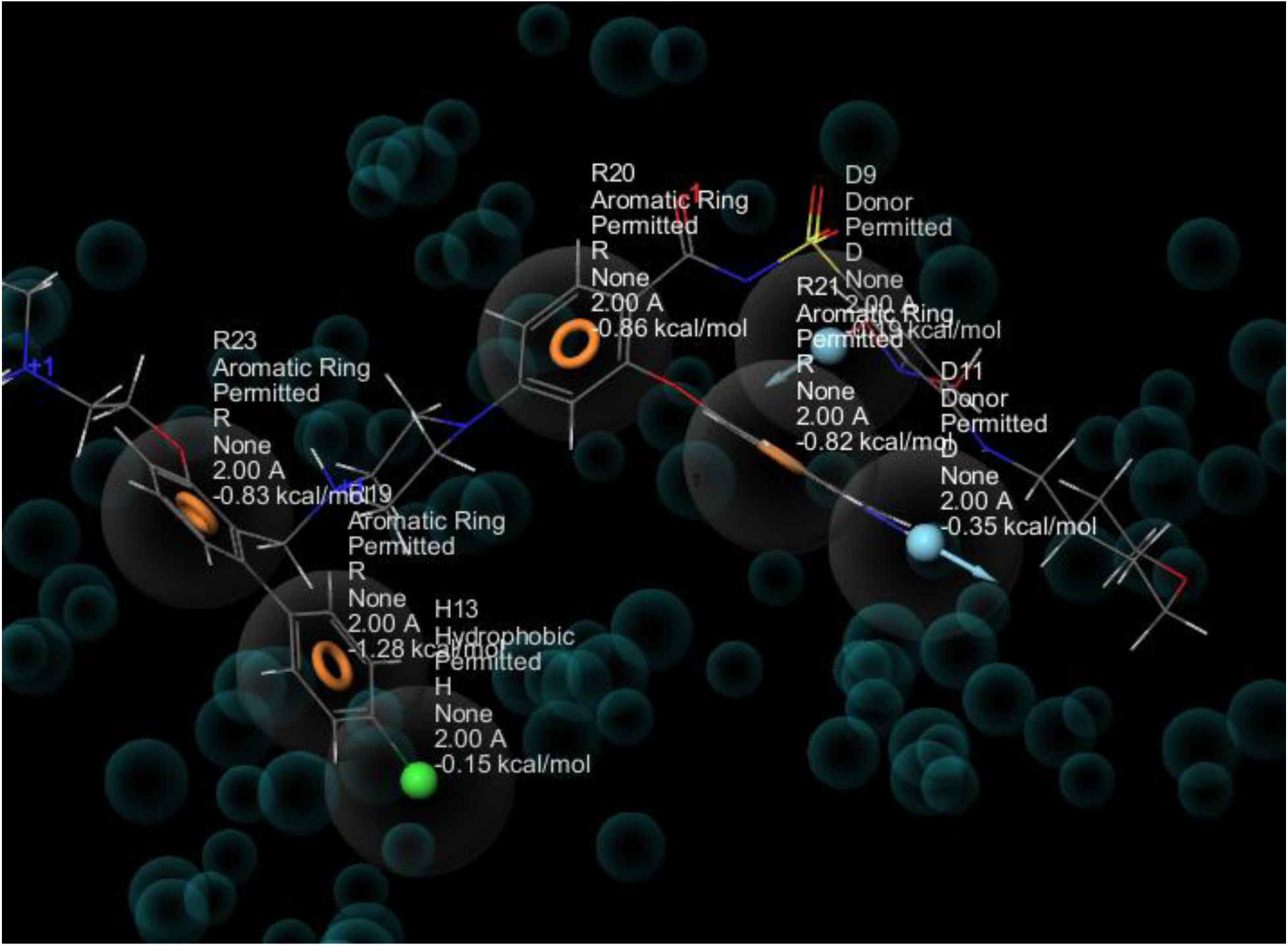
Pharmacophore features of the reference ligand. This features were later used to screen the compounds of *Morus alba,* where a few of them came out with good fitness score. The compounds with favorable fitness scores were then researched as potential inhibitors of Bcl-2.

#### 3.1.1 Fitness score of the top five compounds of *Morus alba*

**Table 1:** Table showing the fitness score of the top five compounds of *Morus alba* after being subjected to the pharmacophore hypothesis generated based on Venetoclax, the reference compound.

| S/N | Compound | Fitness Score |
| --- | --- | --- |
| 1 | **Venetoclax (ABT-199)** | **2.03** |
| 2 | **Navitoclax (ABT-263)** | **1.96** |
| 3 | Quercetin-3-(6-Malonylglucoside) | 2.26 |
| 4 | Epigallocatechin gallate | 2.12 |
| 5 | Quercetin hexoside | 1.86 |
| 6 | Quercetin glucuronide | 1.74 |
| 7 | Mulberroside | 1.64 |

### 3.2 Docking Score

**Table 2:** Table showing the docking score, molecular mechanics with generalized born surface area (MMGBSA) score, and violation of Lipinski’ rule of five violation based on the top five performing compounds of *Morus alba*. The fitness of the compounds were compared to those of the standard drugs, Navitoclax and Venetoclax.

| S/N | Compound | Docking Score (Kcal/mol) | MMGBSA (Kcal/mol) | LROV |
| --- | --- | --- | --- | --- |
| 1 | **Venetoclax (ABT-199)** | **-9.468** | **-68.85** | **3** |
| 2 | **Navitoclax (ABT-263)** | **-9.058** | **-81.83** | **3** |
| 3 | Quercetin-3-(6-Malonylglucoside) | -10.912 | -71.18 | 3 |
| 4 | Epigallocatechin gallate | -9.765 | -57.89 | 2 |
| 5 | Quercetin hexoside | -9.690 | -28.69 | 3 |
| 6 | Quercetin glucuronide | -9.349 | -34.95 | 2 |
| 7 | Mulberroside | - 9.115 | -43.82 | 3 |

### 3.3 ADMETox Parameters

**Table 3:** Table giving some important pharmacokinetic and pharmacodynamics properties of the standard drugs and the test compounds.

| S/N | Compound | QPlogHERG | QPPCaco | QPlogBB | QPPMDCK |
| --- | --- | --- | --- | --- | --- |
| 1 | Venetoclax (ABT-199) | -9.033 | 10.946 | -2.879 | 10.308 |
| 2 | Navitoclax (ABT-263) | -10.008 | 12.86 | -1.465 | 64.131 |
| 3 | Quercetin-3-(6-Malonylglucoside) | -3.009 | 0.399 | -3.957 | 89.13 |
| 4 | Epigallocatechin gallate | -5.065 | 1.581 | -3.835 | 0.464 |
| 5 | Quercetin hexoside | -5.645 | 0.066 | -6.558 | 0.015 |
| 6 | Quercetin glucuronide | -3.18 | 0.46 | -3.727 | 0.155 |
| 7 | Mulberroside | -6.617 | 0.716 | -5.753 | 0.197 |

QPlogHERG: Predicted IC_50_ value for HERG K^+^ Channel Blockage (concern below -5)

QPPCaco: Predicted Caco-2 Cell Permeability. (500 great)

QPPMDCK: Predicted MDCK Cell Permeability. (500 great)

QPlogBB: Predicted brain/blood partition coefficient. –3.0 – 1.2

### 3.4 Interaction Diagram of the Ligands

**Figure 5:**
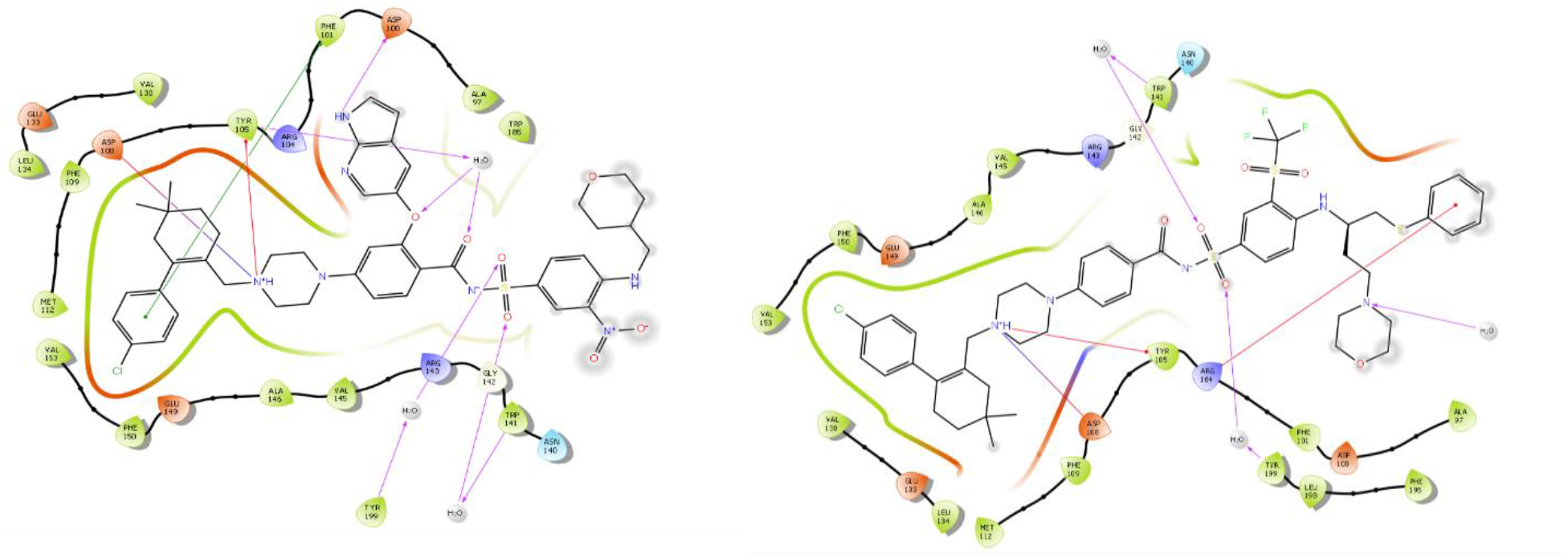
Figure showing the various forms of 2-dimensional interactions between the standard drugs, Venetoclax (left) and Navitoclax (right), and the crystal structure of human B-Cell lymphoma 2 (Bcl-2).

**Figure 6:**
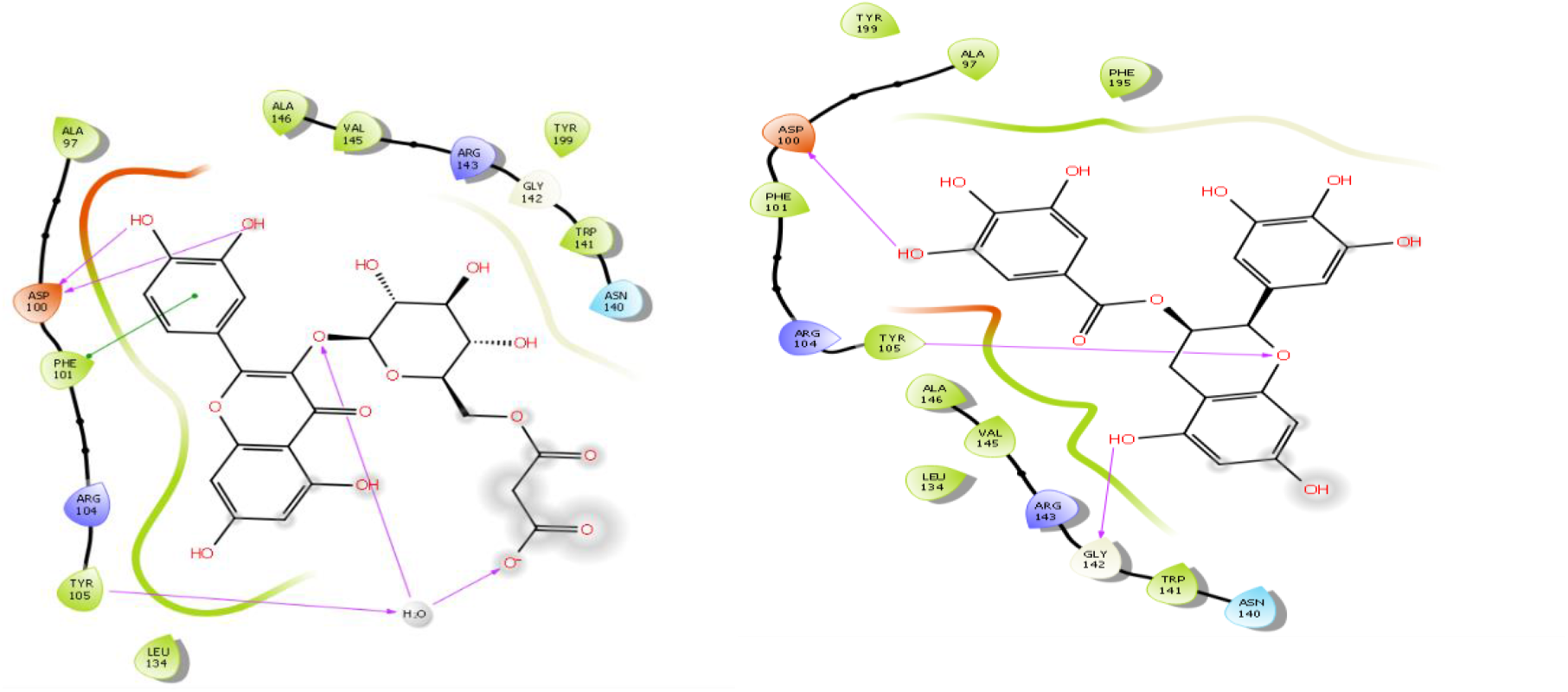
Figure showing the various forms of 2-dimensional interactions between the top two ligands, quercetin-3-(6-Malonylglucoside) (left) and epigallocatechin gallate (right) of *Morus alba* and the crystal structure of human B-Cell lymphoma 2 (Bcl-2).

**Figure 7:**
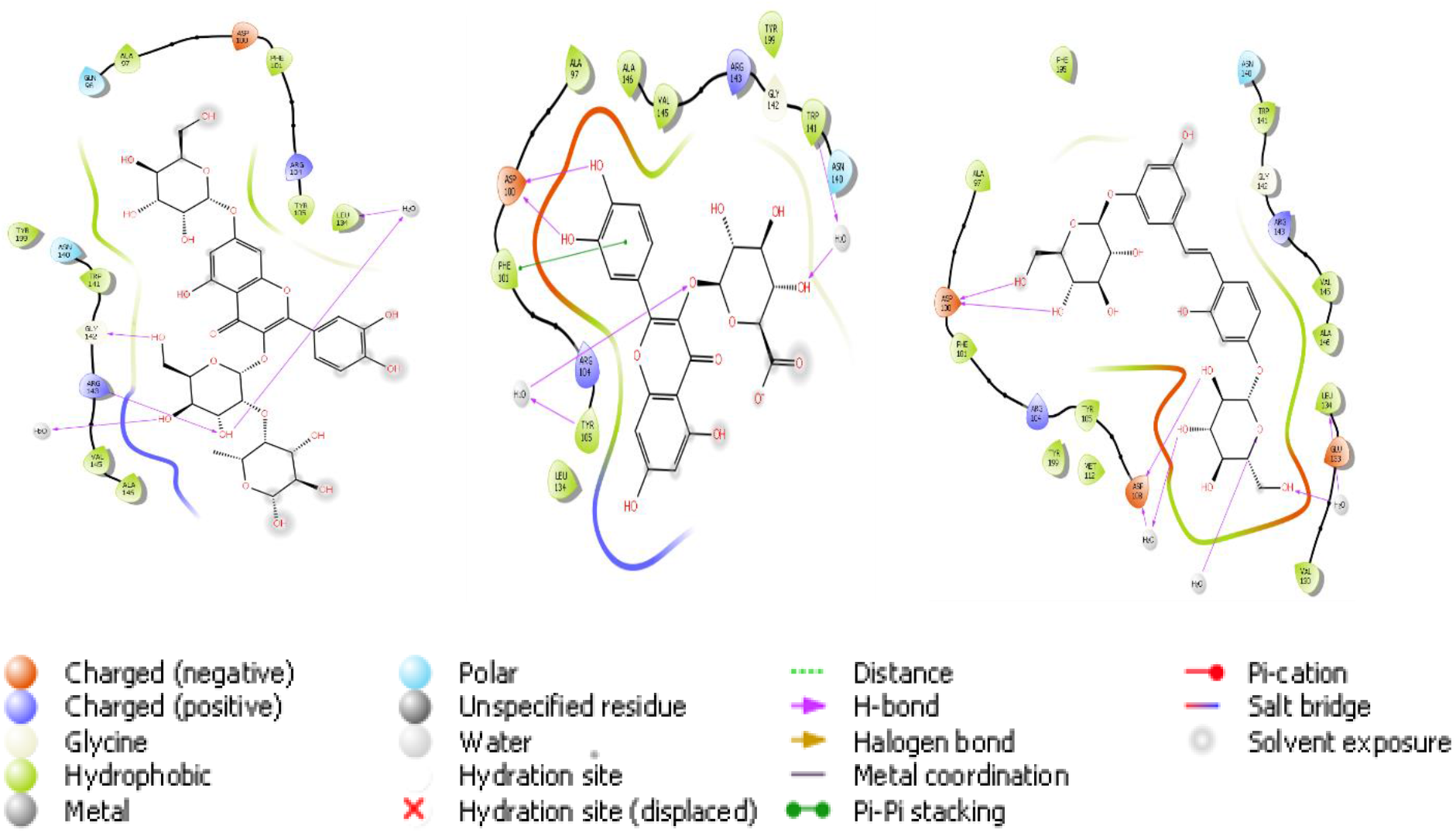
Image of 2-dimensional interaction of Bcl-2 with Quercetin hexoside (left), Quercetin glucuronide (middle), and mulberroside (right). The image gave a summary of the various interactions needed for the stability of the compounds at the active site of the protein.

**Table 4:**
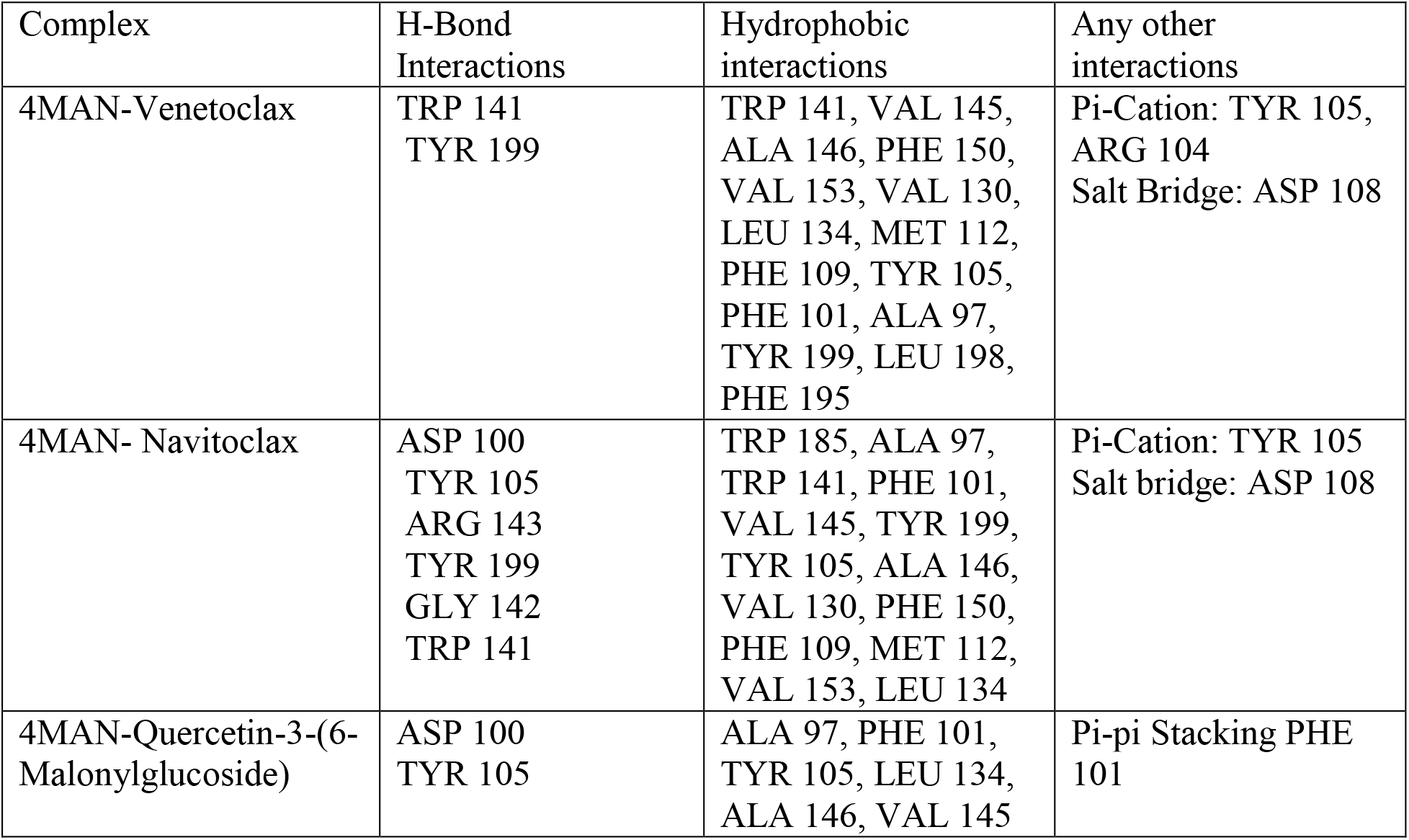
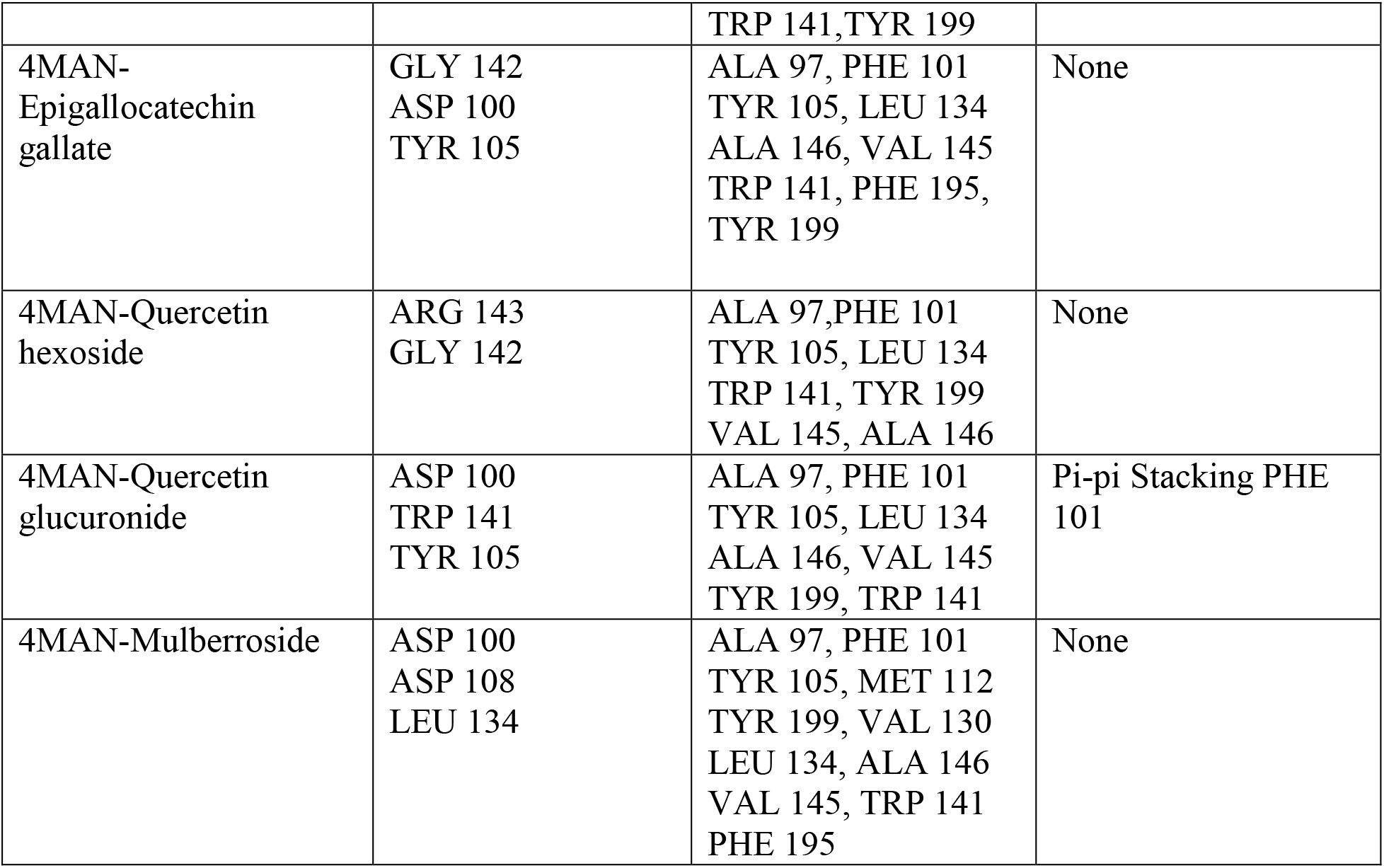
Table giving a summary of the numerous interactions between the ligands and protein. The recorded interactions are crucial to the stability of the compounds at the active site of the protein, Bcl-2, an anti-apoptotic protein.

### 3.5. Global descriptor of reactivity

**Table 5.** Table showing the values of some important global descriptors of drug-like compounds. Important descriptors of reactivity like ionization potential, and electron affinity are used as important parameters for the chemical reactivity and reaction nature of studied biological systems. The ionization potential and electron affinity of a molecule are directly related to HOMO and LUMO values, respectively (Hussain et al., 2021). The various forms of interaction between molecules requires the involvement of electrons.

| Compound | Ionization potential (eV) | Electron affinity (eV) |
| --- | --- | --- |
| Navitoclax | 8.865 | 1.777 |
| Venetoclax | 8.529 | 1.472 |
| Quercetin-3-(6-Malonylglucoside) | 8.911 | 0.620 |
| Epigallocatechin gallate | 8.993 | 0.696 |

**Figure 8:**
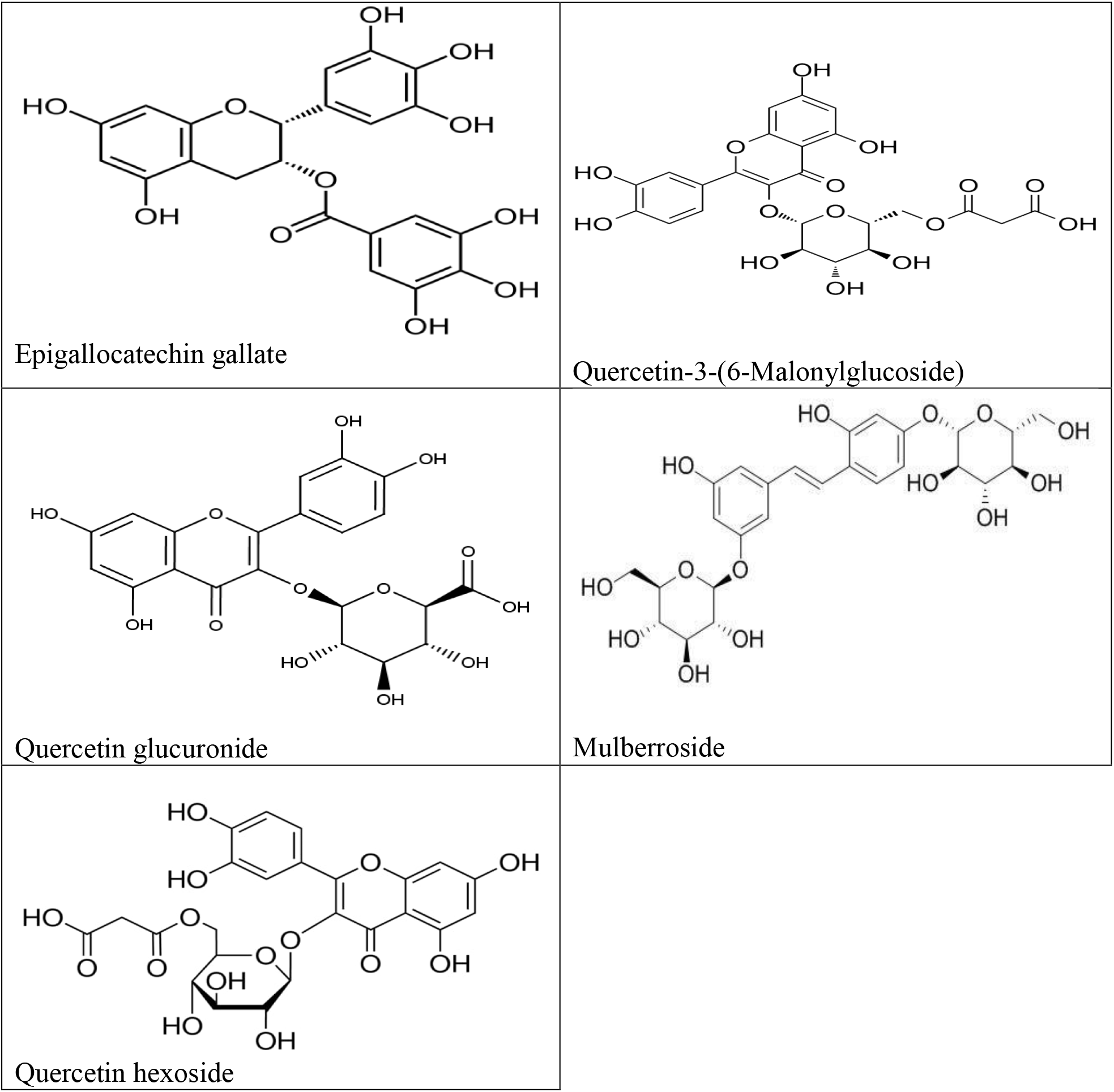
Images of the top five compounds reported in *Morus alba*.

**Figure 9:**
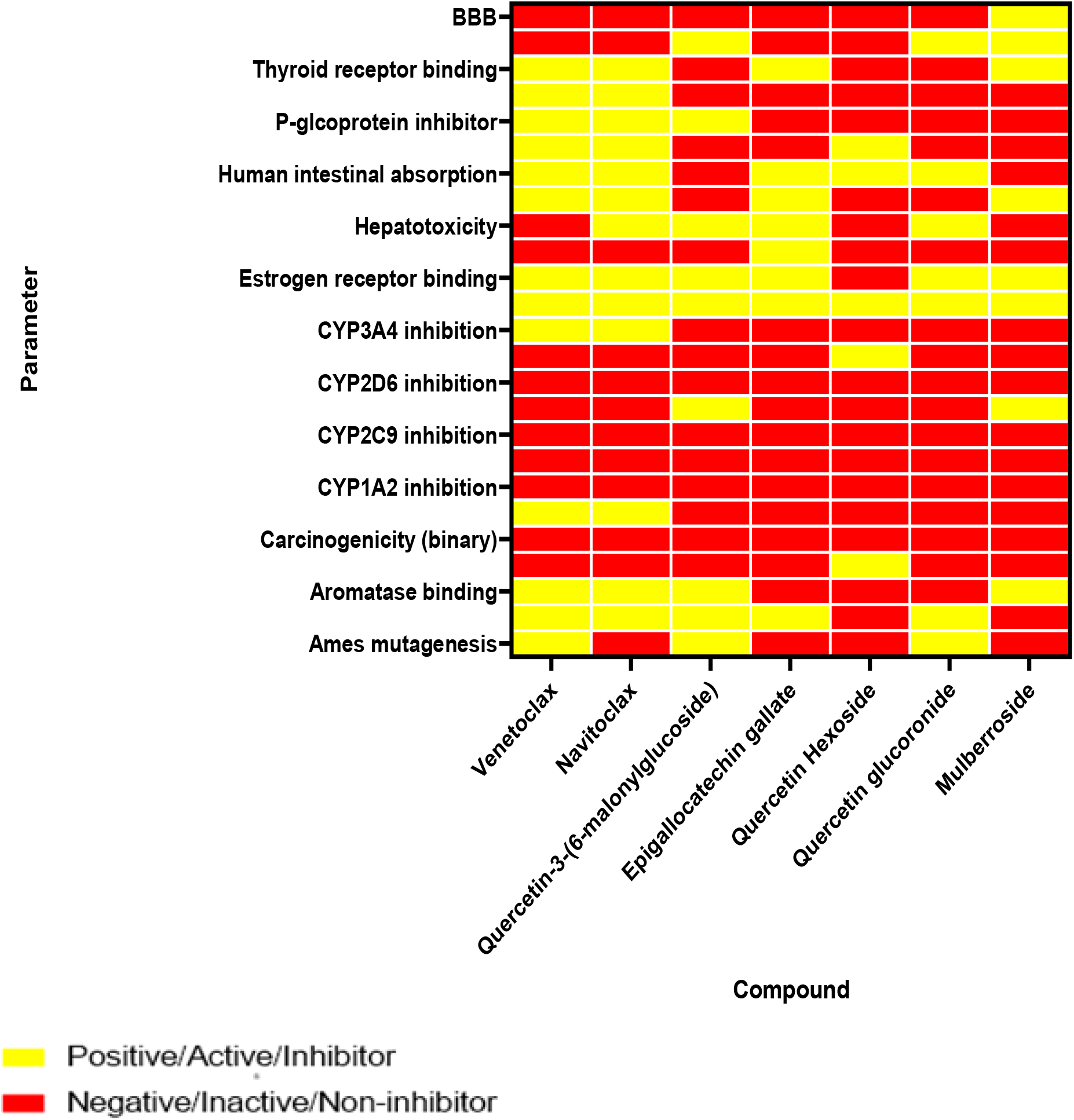
Heat Map showing some pharmacokinetic and pharmacodynamics properties of the standard drugs and the top five ligands of *Morus alba*.

**Figure 10:**
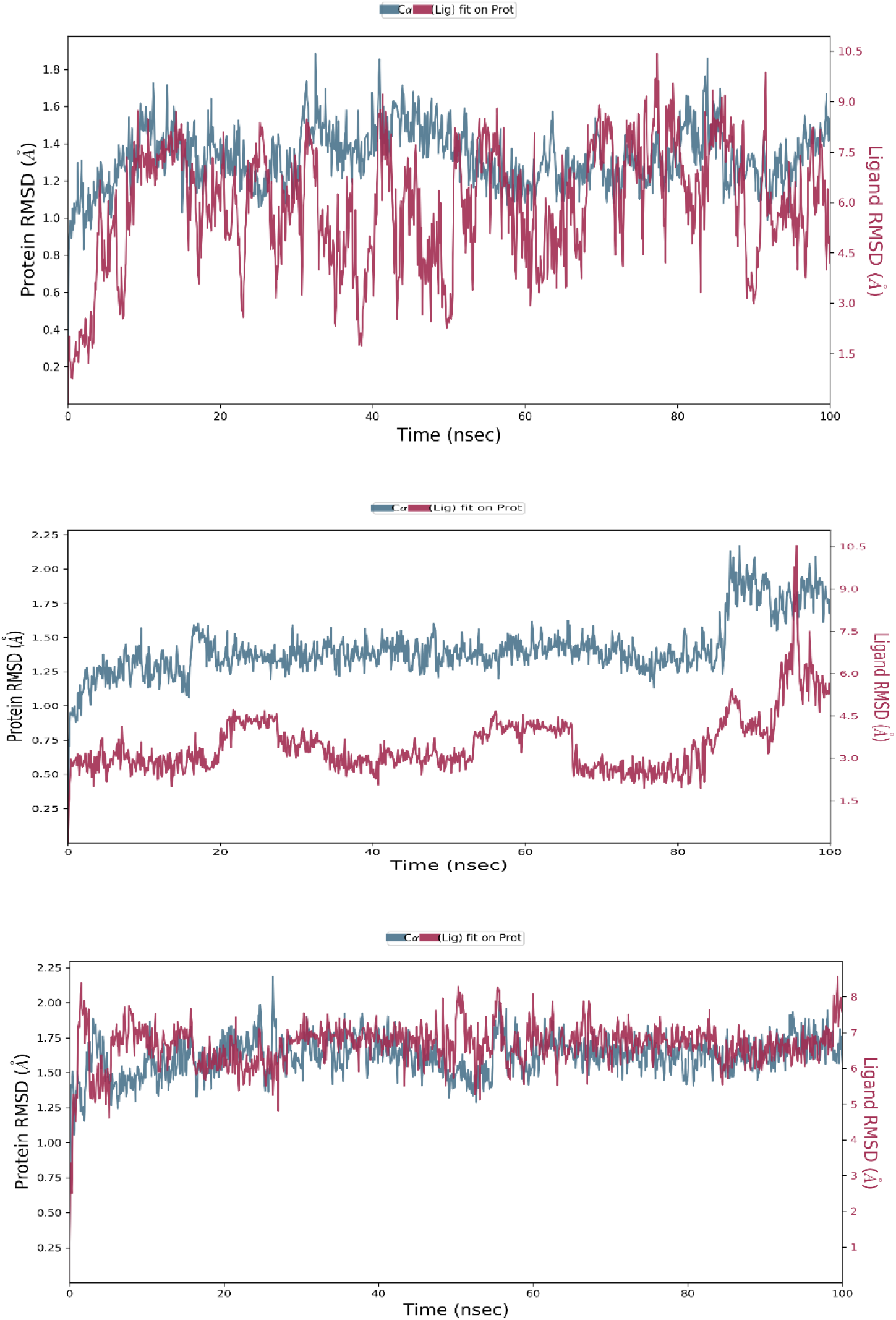
Protein-ligand RMSD plots for 4MAN-Venetoclax (top), 4MAN-Navitoclax (middle), and 4MAN-Quercetin-3-(6-malonylglucoside) (down) complexes. The blue spectrum represents the alpha carbon of the backbone of the protein with its scale on the left of the y-axis while the red spectrum represents the ligands with its scale on the right of the y-axis. All graphs have various scales; the y-axis of the scales is in angstrom (Å) while the x axis is in nanoseconds (nsec).

**Figure 11:**
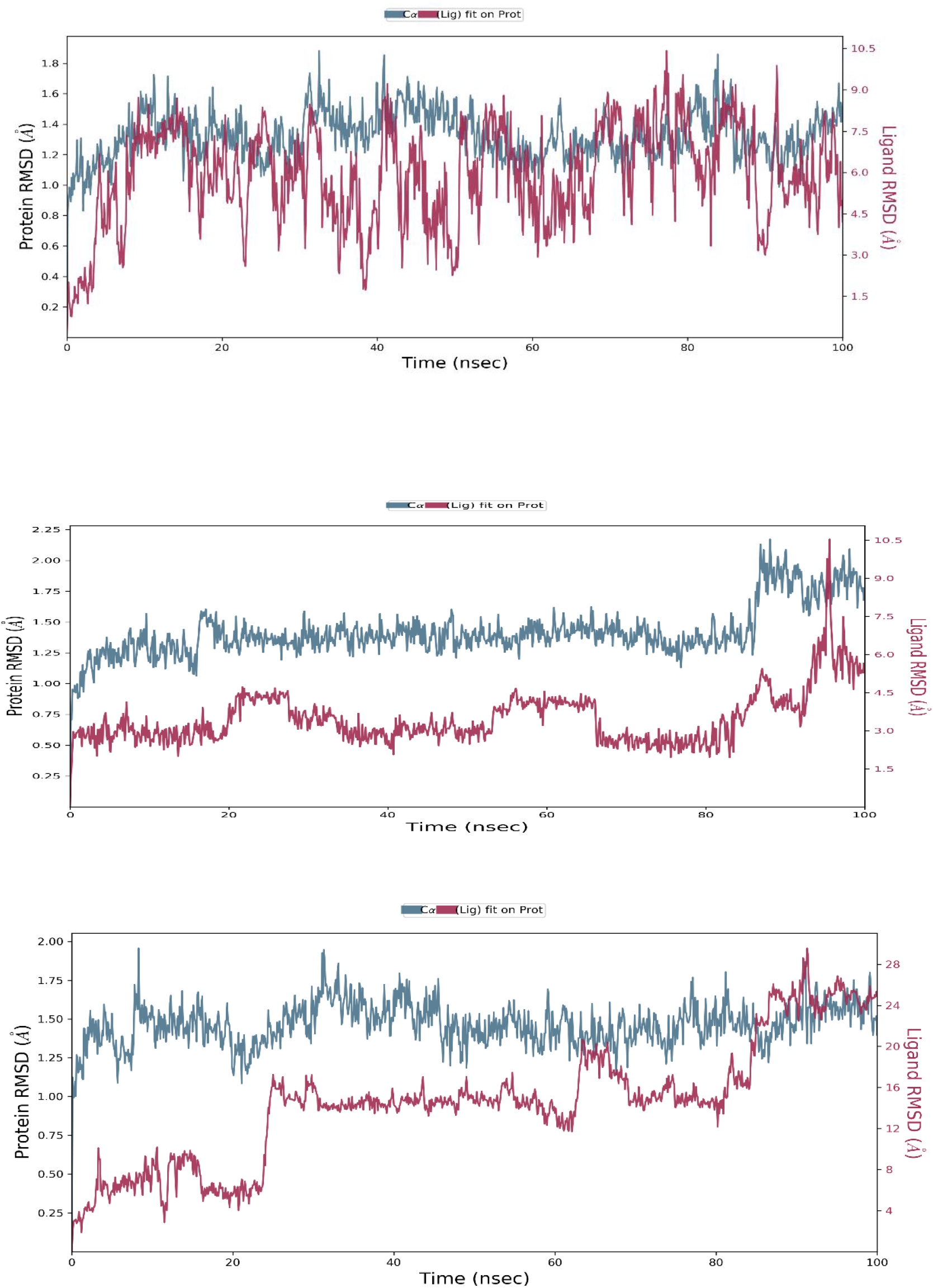
Protein-ligand RMSD plots for 4MAN-Venetoclax (top), 4MAN-Navitoclax (middle), and 4MAN-Epigallocatechin gallate (down) complexes. The blue spectrum represents the alpha carbon of the backbone of the protein with its scale on the left of the y-axis while the red spectrum represents the ligands with its scale on the right of the y-axis. All graphs have various scales; the y-axis of the scales is in angstrom (Å) while the x axis is in nanoseconds (nsec).

**Figure 12.**
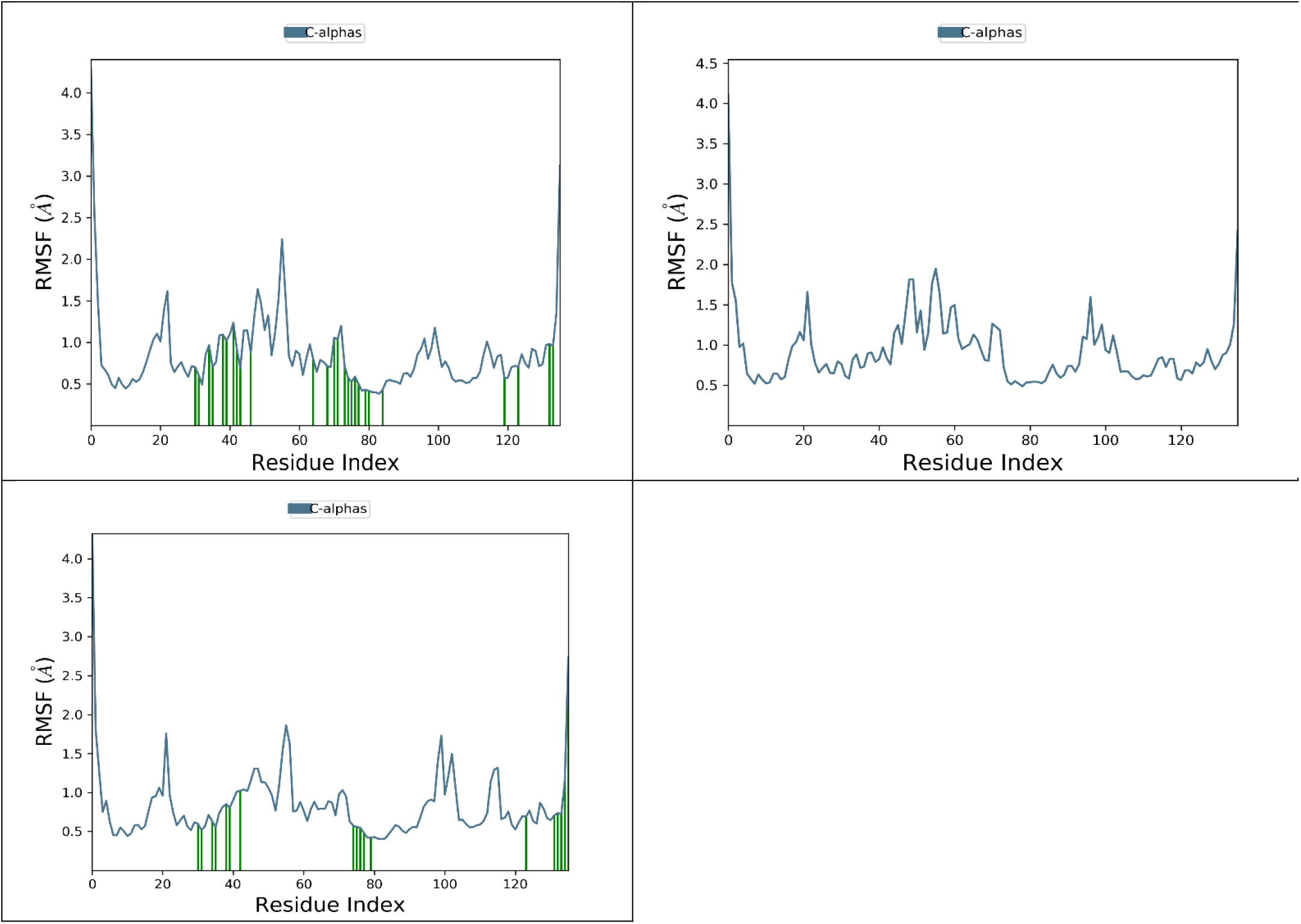
Protein-Ligand Root Mean Square Fluctuation (RMSF). 4MAN-Venetoclax (top left), 4MAN-Navitoclax (top right), and 4MAN-Quercetin-3-(6-malonylglucoside) (down) complexes. The blue lines represent the alpha carbon of the backbone of the protein, while the green bar represents the ligands making contact with the protein.

**Figure 13.**
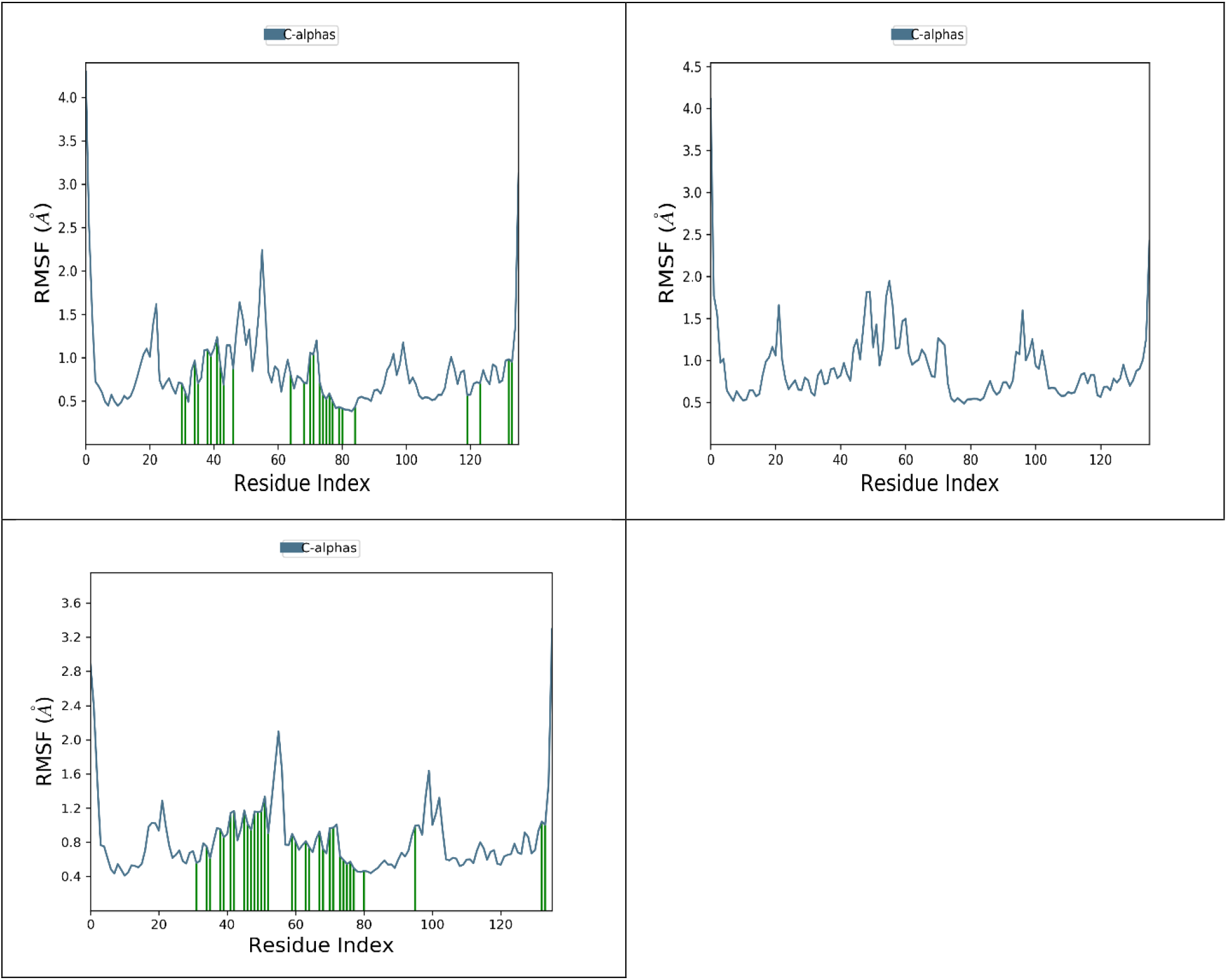
Protein-Ligand Root Mean Square Fluctuation (RMSF). 4MAN-Venetoclax (top left), 4MAN-Navitoclax (top right), and 4MAN-Epigallocatechin gallate (down) complexes. The blue lines represent the alpha carbon of the backbone of the protein, while the green bar represents the ligands making contact with the protein.

**Figure 14.**
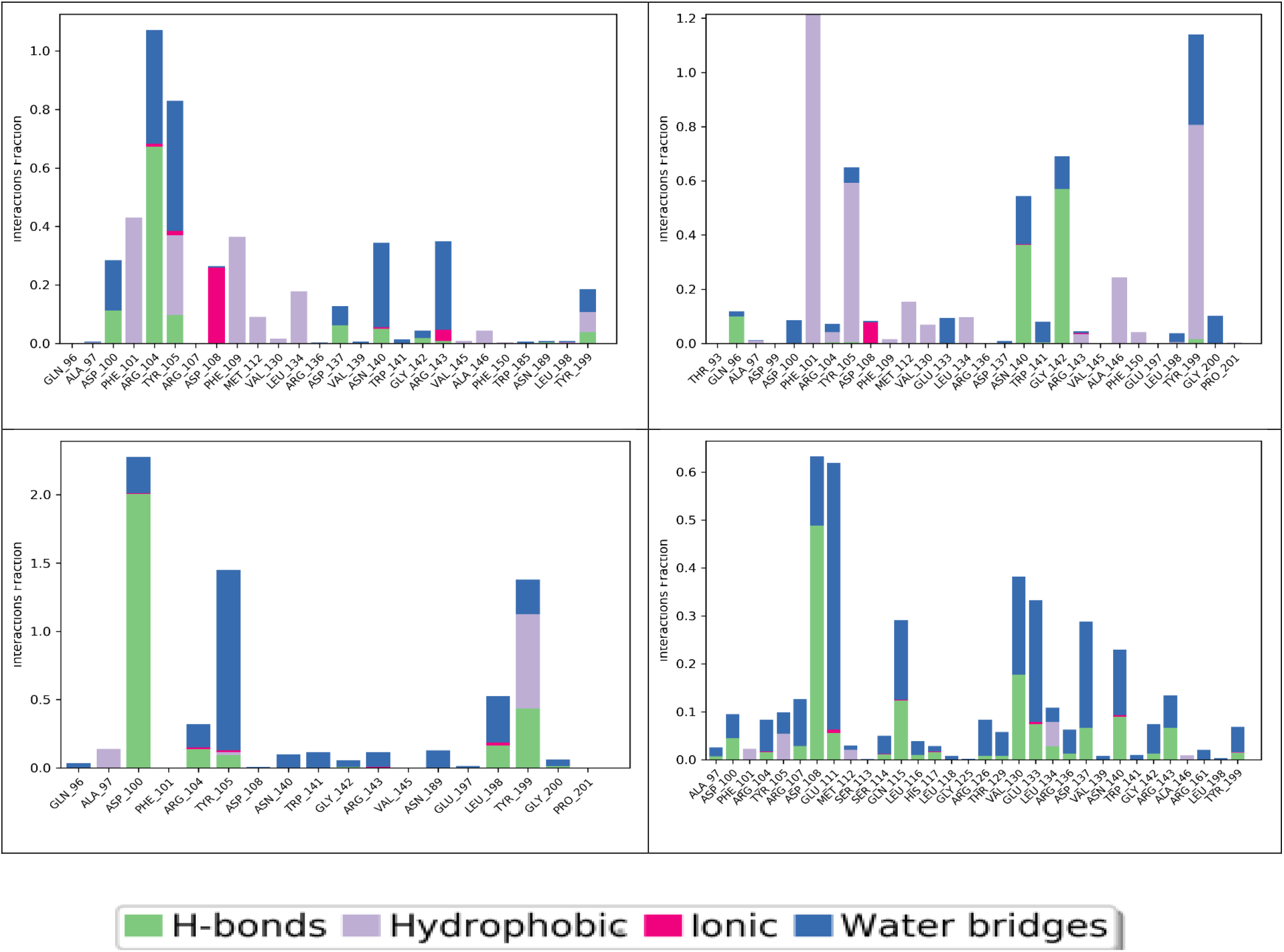
Molecular interaction between 4MAN-Venetoclax (top left), 4MAN-Navitoclax (top right), 4MAN-Quercetin-3-(6-malonylglucoside) (bottom right), and 4MAN-Epigallocatechin gallate (bottom right) complexes. The Y axis represents the interactions fraction scale, while the x axis represents the positions of amino acids in the protein.

**Figure 15.**
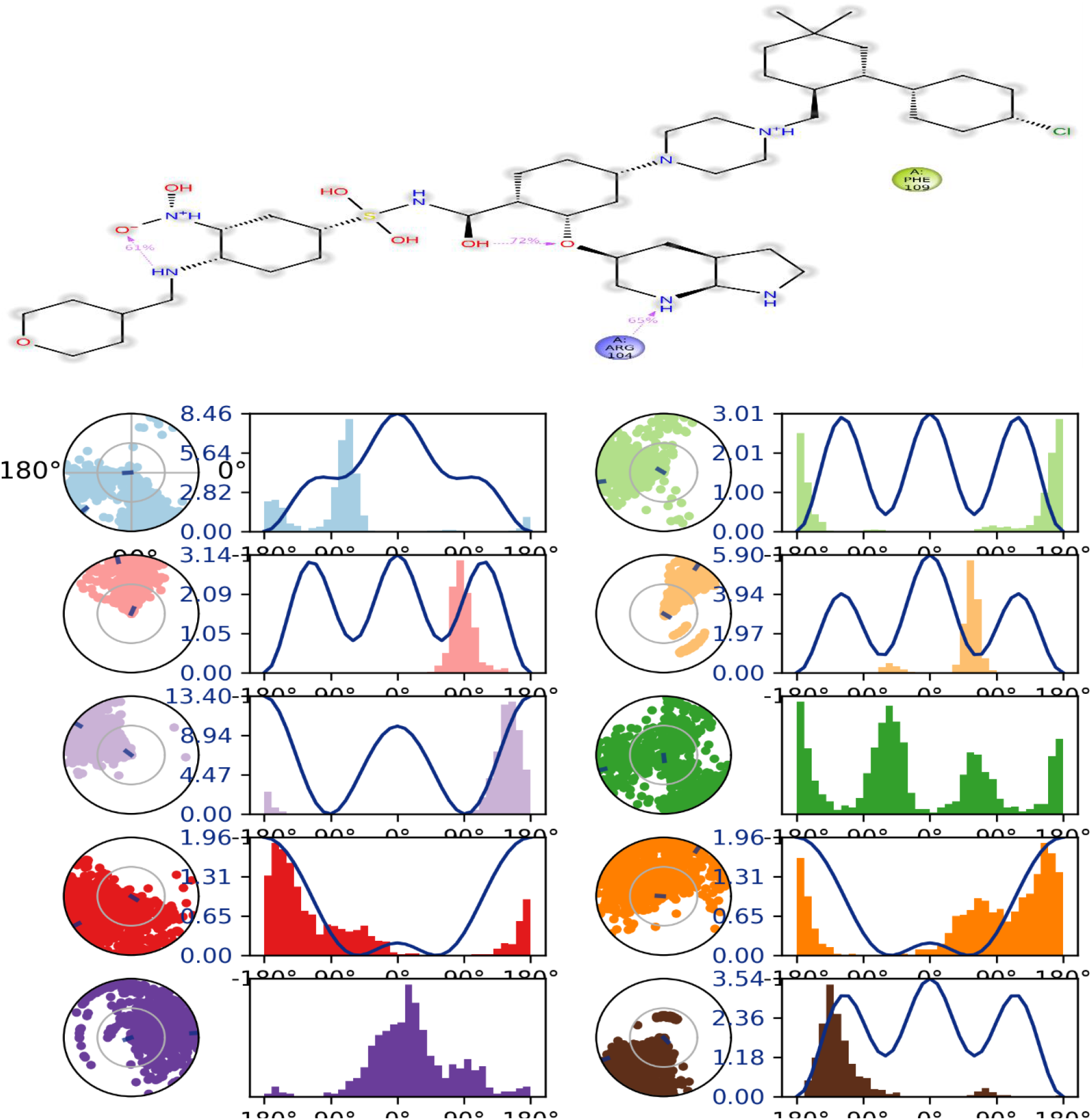
Ligand torsion profile of the standard drug, Venetoclax. Each rotatable bond in its 2D structure is encoded with a color accompanied by a radial plot and a bar plot with the same color coding. The dial (or radial) graphs show the simulation torsional conformation.

**Figure 16.**
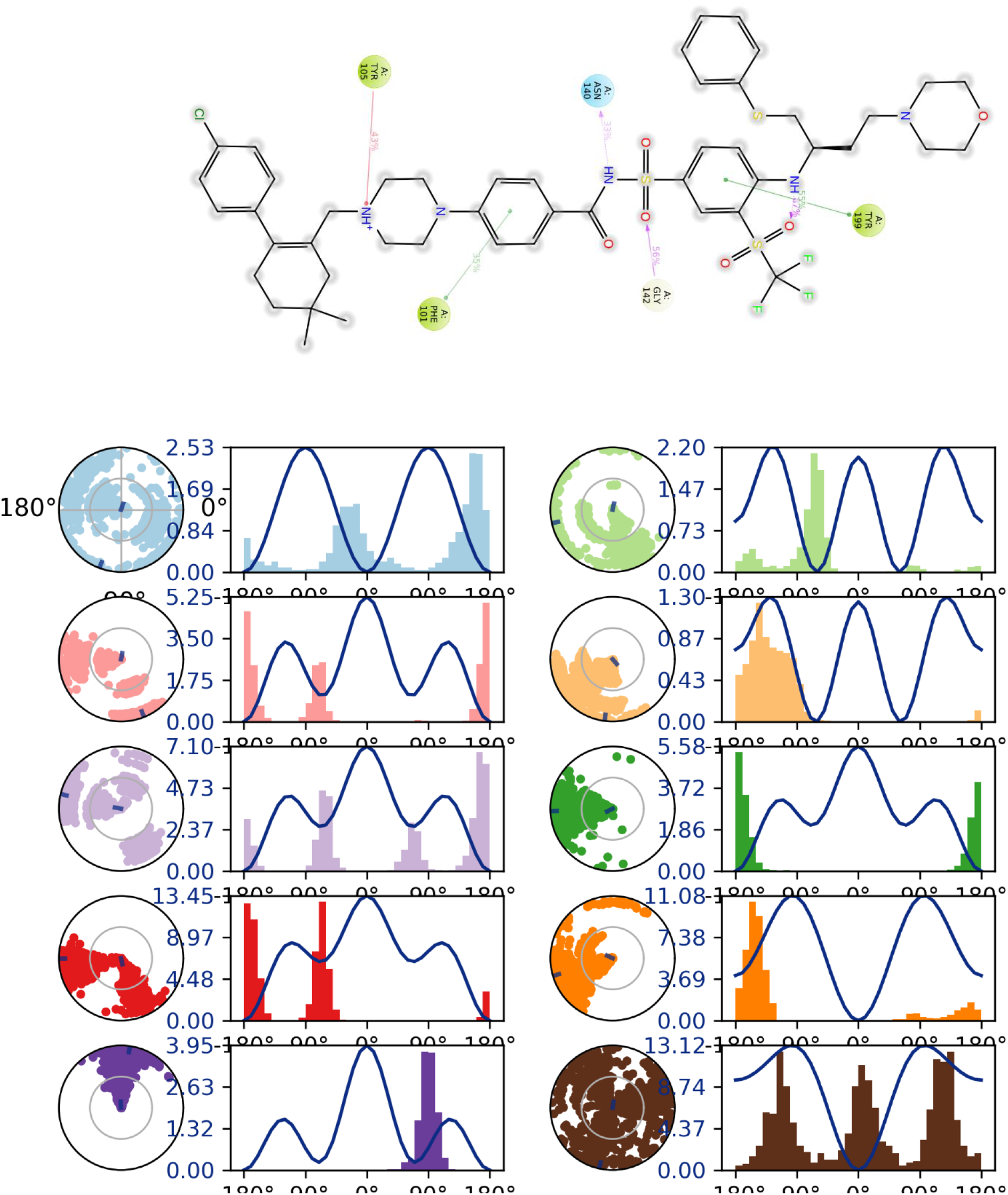
Ligand torsion profile of the standard drug, Navitoclax. Each rotatable bond in its 2D structure is encoded with a color accompanied by a radial plot and a bar plot with the same color coding. The dial (or radial) graphs show the simulation torsional conformation.

**Figure 17.**
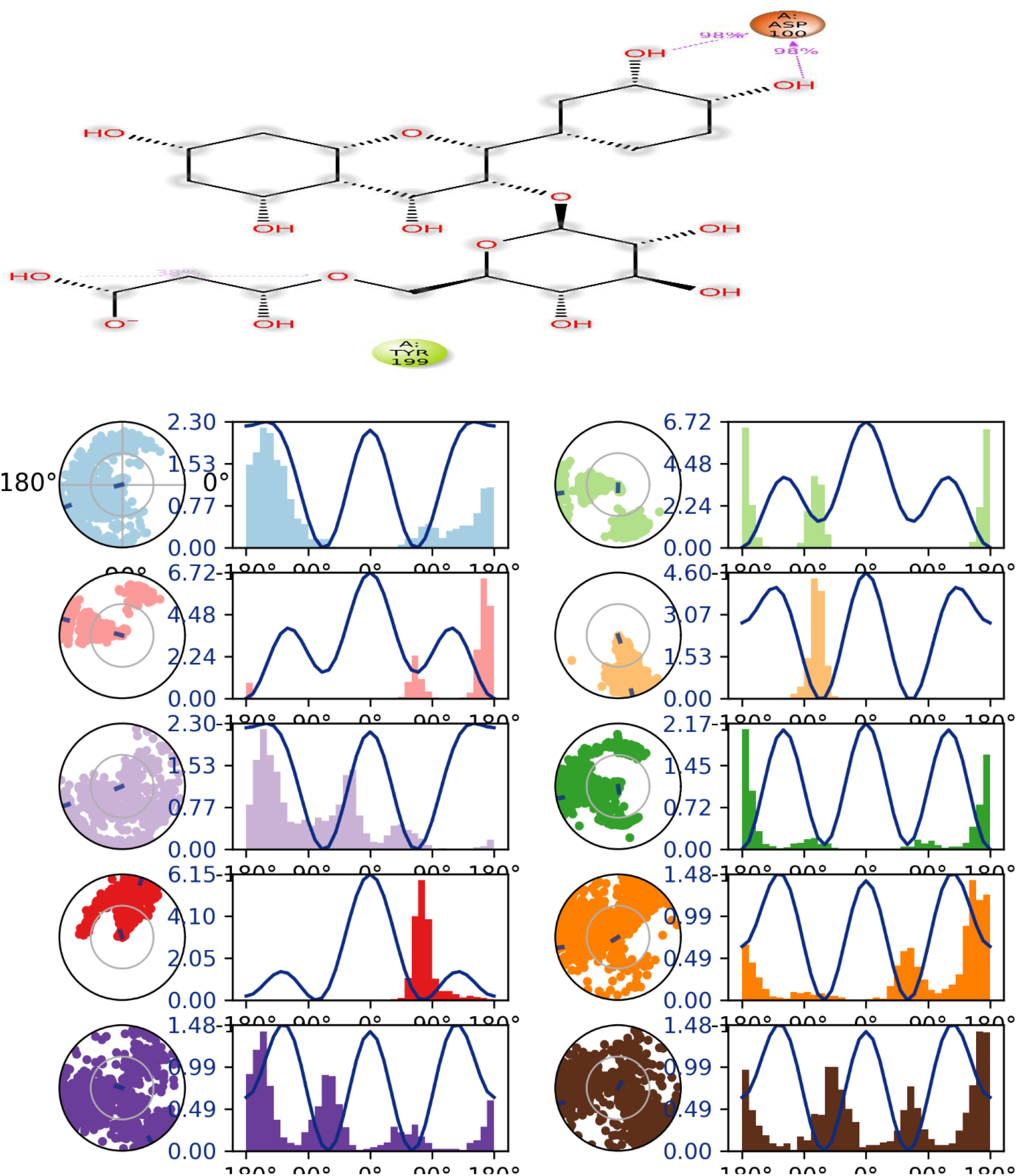
Ligand torsion profile of the topmost test compound, Quercetin-3-(6-Malonylglucoside). Each rotatable bond in its 2D structure is encoded with a color accompanied by a radial plot and a bar plot with the same color coding. The dial (or radial) graphs show the simulation torsional conformation.

**Figure 18.**
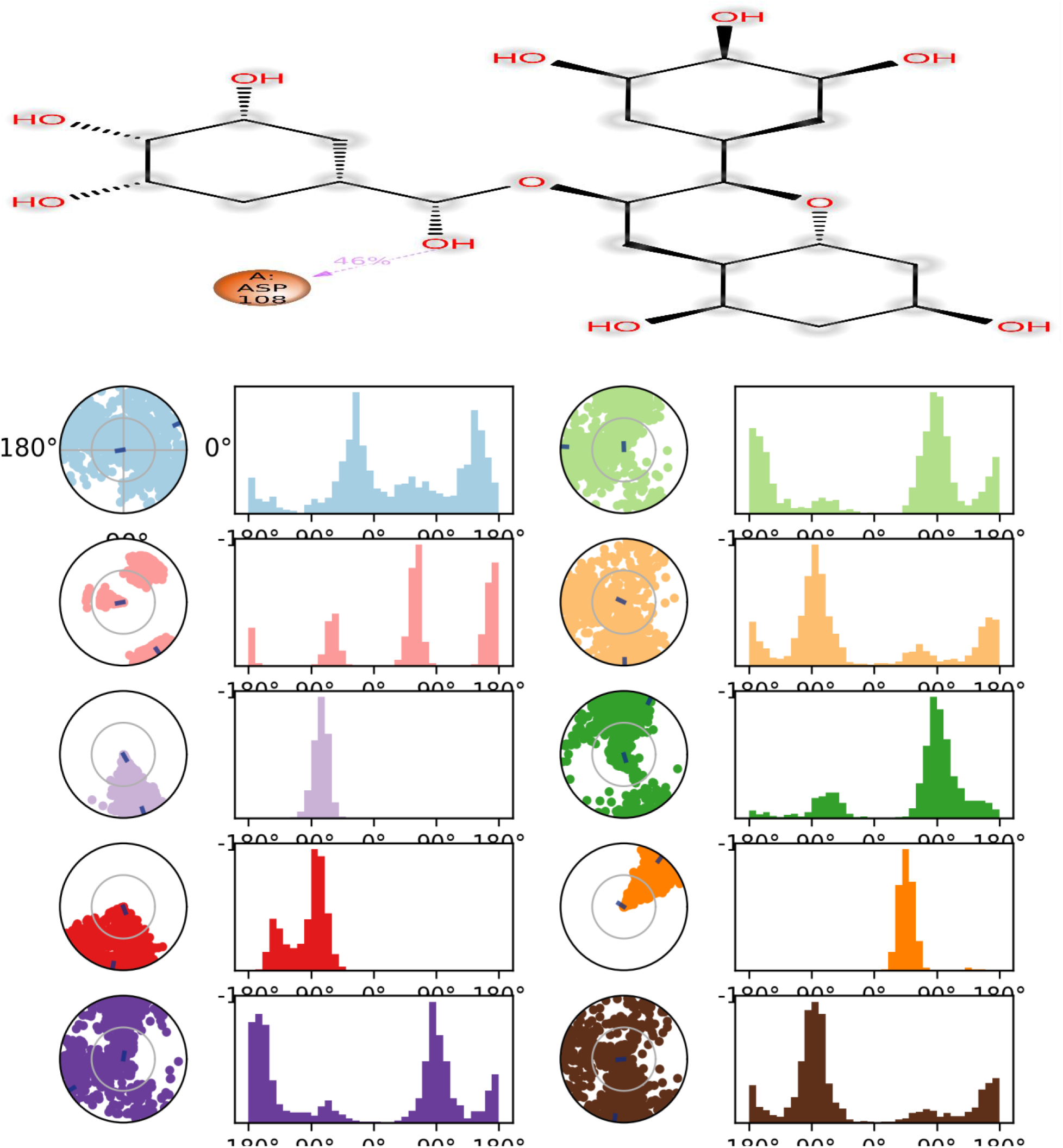
Ligand torsion profile of the second ranked compound, Epigallocatechin gallate. Each rotatable bond in its 2D structure is encoded with a color accompanied by a radial plot and a bar plot with the same color coding. The dial (or radial) graphs show the simulation torsional conformation.

## 4.0 Discussion

### 4.1 Pharmacophore, Molecular Docking, MM-GBSA, LRoV, and ADMET.

A pharmacophore represents the three-dimensional model of key features that are responsible for specific biological activities of chemical entities (Van Drie, 2003). These features include hydrogen bond donors, acceptors, and hydrophobic regions, which provide structural basis for molecule-protein interactions. The pharmacophore hypothesis was set based on the interactions of the standard drug, Venetoclax, in the binding site of the target protein. This helped to identify lead compounds with the expected features for a better binding affinity. The fitness of the top five compounds of mulberry were then compared with those of the standard drugs used in this work, and the result presented in (table 1). Molecular docking techniques were then used to simulate the interaction between the compounds and the target protein. Molecular docking represents a computational technique used in drug discovery and molecular biology research to predict the binding orientation and affinity of small molecules (ligands) with target protein (s) (Bodun et al., 2023). The technique plays a crucial role in understanding the molecular interactions between drugs and their protein targets. Its result, called docking score, is a quantitative measure used to evaluate the quality of the predicted binding modes and guide further experimental studies. From table 2, the binding affinity of the standard drugs were compared with those of the selected lead compounds from the plant. Here in table 2, the molecular mechanics with generalized born surface area (MM-GBSA) results were presented. MM-GBSA plays a significant role in drug discovery by providing valuable insights into the binding affinities of small molecules to target proteins. MM-GBSA calculations can estimate the binding affinity between a small molecule (ligand) and its target protein. By calculating the free energy difference between different ligand-protein complexes, MM-GBSA can rank and prioritize potential drug candidates based on their predicted binding affinities. This information aids in lead optimization and selecting compounds with the highest likelihood of binding to the target effectively (Wu et al., 2021). The MM-GBSA studies indicated that the compounds are bound favorably to the active site of the target protein, as indicated by the values presented in table 2. The Lipinski’s rule of five (LRoV) also indicated that the lead compounds compared favorably with the standard drugs used in the inhibition of anti-apoptotic Bcl-2. The Lipinski rule of five suggests that compounds meeting the criteria in the rule (>500Da molecular weight, >140Å topological surface are, ≥5 H-bond donor, ≥10 H-bond acceptor, and >5 log P value) are more likely to have favorable pharmacokinetic properties, including good oral absorption, distribution, and bioavailability (Chen et al., 2020). However, it is important to note that the rule is not an absolute predictor of a compound’s success as a drug but serves as a rough guideline during the early stages of drug discovery (Segall, 2012). The standard drugs each violated three of the five rules, a feature repeated by some of the lead compounds for example, quercetin-3-(6-malonylglucoside), but was bettered by some, e.g. epigallocatechin gallate as reported in table 2. The pharmacokinetic properties of the lead compounds were also assessed and presented in table 3. From the table, the predicted IC_50_ for the HERG K+ channel for the standard drugs were compared with those of the test compounds. The human ether-a-go-go-related gene potassium (HERG K+) channel is an ion channel protein that plays a crucial role in cardiac repolarization (Perrin et al., 2008). It is a member of the voltage-gated potassium channel family. Dysfunction or impairment of the HERG channel can lead to abnormalities in cardiac repolarization, potentially resulting in a condition known as long QT syndrome (LQTS). The predicted Caco-2 cell permeability, Madin-Darby Canine Kidney (MDCK) cell permeability, and brain/blood partition coefficient were also reported in table 3. Furthermore, global reactivity in terms of ionization potential and electron affinity were also compared between the standard drugs and top compounds, and the result of the comparison was summarized in table 5. For biological effects, ligand and protein molecules must interact. Whatever form of interaction that occur, it must involve electrons. Since ionization potential and electron affinity measure the amount of energy needed to lose and acquire electron respectively, they are therefore important in drug discovery. In drug discovery, this concept (ionization potential and electron affinity) is extended to the energy required for drug molecules to interact with biological membranes, affecting their permeability and ability to cross cellular barriers.

### 4.2 Molecular dynamics simulation

Molecular dynamics (MD) simulation of biomolecular systems has expanded to study a wide variety of important biological processes. Due to interdisciplinary advancements in biological sciences and the increased use of tools of bioinformatics, the screening of synthetic as well as natural compounds has revolutionized the field of drug discovery. One of the main strengths of using molecular dynamics simulation in drug discovery is that it enables detailed analysis of the interaction between protein and ligand at the molecular level, revealing potential drug targets and aiding in the design of new drugs (Omoboyowa et al., 2022). Another significant strength of the MD simulation is its ability to simulate protein and protein-ligand interactions at millisecond timescales with high resolution. This enables scientists to study the stability and dynamics of complex biomolecular systems, including their conformational changes and interactions (Elkaeed et al., 2022). MD simulation was therefore used to study the various interactions among the ligands and protein used in this work. This was done to reduce and possibly eliminate completely any level of doubt about the potency of the ligands inhibiting Bcl-2 at molecular level.

#### 4.2.1 Root mean square deviation (RMSD)

In computational biology, RMSD is a technique that is widely used to assess how much macromolecules veer over time (Sargsyan et al., 2017). In this work, we examined the structural stability of Venetoclax, Navitoclax, Quercetin-3-(6-Malonylglucoside), and Epigallocatechin gallate at 100ns simulation timescale. Venetoclax and Navitoclax were chosen as standards for the simulation because they have gained much attention as potent inhibitors of Bcl-2. The behavior of the compounds (standards and top two of *Morus alba*) in the hydrated environment of the Bcl-2 active sites were evaluated during the simulation run. Before calculating the RMSD based on the atom selection, in this case on the alpha Carbon, all protein frames are first aligned on the reference frame backbone. How stable a ligand is in relation to a protein and its binding pocket is indicated by the ligand RMSD (Allegra et al., 2021). Since it has been stated that changes of the order of 1-3 Å are perfectly acceptable for small, globular proteins, and that changes much larger than that, however, mean that such protein is undergoing a large conformational change during the simulation, figures 9 and 11 show that the RMSD values of the two top compounds fall within the permissible range, and that the system is stable during the period of simulation. However, the RMSD of the 4MAN-Quercetin-3-(6-malonylglucoside) complex showed to be better than that of the second ranked compound of *Morus alba*, with this favoring the likely use of Quercetin-3-(6-Malonylglucoside) as potent drug compound over epigallocatechin gallate. As a corollary, it is important that the simulation converges, meaning that the RMSD values stabilize around a fixed value. If the RMSD of the protein is still increasing or decreasing on average at the end of the simulation, then the system has not equilibrated, and the simulation may not be long enough for rigorous analysis. Fortunately, in this case, the RMSD values of the complexes attained stability towards the end of the simulations, pointing to the fact that the systems have equilibrated, that is, the values are fit for rigorous analysis. Overall, the two best compounds from mulberry that were subjected to simulation showed stability with little fluctuation falling within permissible range throughout the period of simulation. Ligand RMSD (presented on right Y-axis of figures 9 and 11) shows how stable the ligand is with respect to the protein and its binding pocket. From the two figures (10 and 11), ’Lig fit Prot’ shows the RMSD of a ligand when the protein-ligand complex is first aligned on the protein backbone of the reference and then the RMSD of the ligand heavy atoms is measured. As a rule, if the values observed are significantly larger than the RMSD of the protein, then it is assumed that the ligand has diffused away from its initial binding site. With respect to Quercetin-3-(6-Malonylglucoside), there is negligible difference between the protein and ligand RMSD, indicating that the ligand has not diffused away from its original position in the course of the simulation. On the other hand, there seem to be an appreciable difference between the protein RMSD and Epigallocatechin gallate RMSD, a likely pointer to the fact that the ligand has diffused away from its initial position. In any case, such observation is not self-sufficient in preventing the ligand from being used as potent inhibitor of Bcl-2.

#### 4.2.2 Root mean square fluctuation (RMSF)

The RMSF is useful for characterizing local changes along the protein chain (Boyenle et al., 2022). Figures 12 and 13 show the RMSF of the residues at the active site of the protein (4MAN) complexed with Venetoclax, Navitoclax, Quercetin-3-(6-Malonylglucoside), and Epigallocatechin gallate. From the images, virtually all the compounds showed identical behaviors at the catalytic residues of the protein in that they display similarity in lowest fluctuation value (0.5Å). However, Quercetin-3-(6-Malonylglucoside) showed the least value for fluctuation peak (1.8Å), indicating that there is a very strong grip on the ligand by the catalytic residues in the protein. Amino acids that form a protein’s catalytic site should not vary greatly. This explains the fact that the proteins’ active areas are intact enough to accept the inhibitor (Boyenle et al., 2022). Overall, most of the large fluctuations in proteins occur in the loop regions because secondary structure such as alpha helices and beta strands are often more rigid (Daood et al., 2020).

#### 4.2.3 Intermolecular interactions

Protein-ligand interactions have been reported to play critical roles in various biological processes, including signal transduction, metabolism, and gene regulation. To have a clearer picture of the mechanism of protein-ligand interactions, it is important to identify the forces that are responsible for the affinity and specificity between ligands and proteins (Jayaraman et al., 2021). Amongst the various intermolecular interactions that occur during the protein-ligand interaction, hydrophobic and hydrogen bonds have been studied extensively. Numerous studies have reported the importance of both hydrophobic and hydrogen bonds in stabilizing protein-ligand complexes. However, the question of which interaction is stronger in protein-ligand interactions has been widely debated. Recently, research suggests that hydrophobic interactions are stronger than hydrogen bonds in ligand-protein interactions (Patil et al., 2010). Moreover, studies have reported that hydrophobic interactions provide valuable strength for structurally complex stabilization, thus increasing the affinity of the ligand toward the protein (Ahmed et al., 2020). From the work presented herein, the various forms of interaction between the protein and ligands were shown in figures 5, 6, and 7, and later summarized in table 4. Molecular dynamics simulation further validated these interactions earlier seen with docking as depicted by figure 14.

#### 4.4.4 Ligand Torsion

The ligand torsional illustrations for the standards and test compounds were displayed in figures 15, 16, 17, and 18. These images, when carefully studied, give information on which conformation of the compounds is most stable and fit to be used as inhibitor for Bcl-2. In drug discovery, ligand torsional diagrams can aid in the optimization of pharmacophores. As described earlier, pharmacophores are the essential structural and chemical features of a ligand that are responsible for its biological activity. By studying the torsional diagram, researchers can analyze the conformational preferences of ligands and identify the most favorable conformations that interact effectively with the target protein.

### 4.5 Conclusions

*Morus alba* (mulberry) has a rich record of use for both nutritional and therapeutic purposes. Recently, many studies have focused on the identification of potent and less toxic inhibitors from natural sources (plants, animals, etc.), and some of these inhibitors have found their way to the market, while some still remain in clinical trials. Our study, without doubts, concludes that compounds of mulberry, particularly the top two reported here, will be effective compounds in inhibiting anti-apoptotic Bcl-2, and that these compounds could be better optimized as potent inhibitors of Bcl-2 in cancer treatment. Therefore, compounds of mulberry should be subjected to further experimental studies (*in vitro* and *in vivo)* in order to confirm the findings reported herein.

## Acknowledgment

The authors are deeply grateful for the help, encouragement, and material accessibility given by Professor O. I. Omotuyi, Molecular Biology & Simulations (Mols & Sims) Centre.

## Data Availability Statement

The corresponding author will provide access to all docking structures and other documents upon reasonable request.

## Author Contributions Statement

E. S. Omirin, E. A. Olugbogi, O. I. Omotuyi, O. G. Afokhume, and M. A. Aderiye concived and developed the idea behind the work; E. F. Okoh and S. O. Boboye took care of the MD simulations; E. S. Omirin, O. O. Agosile, O. O. Adelegan, and B. O. Ibitoye did the writing and proof-reading of the manuscript.

